# Modulation of mitochondria-ER contacts decrease inflammasome formation and restores amyloid β-peptide phagocytosis in adult mouse microglia

**DOI:** 10.64898/2026.03.10.710773

**Authors:** Ming Ho Choi, Luana Naia, Maria Ankarcrona

## Abstract

**Background:** Alzheimer’s disease (AD) is the most prevalent neurodegenerative disease, currently devoid of a cure. AD’s clinical manifestations stem from a multitude of dysfunctional cellular processes, regulated by mitochondria-endoplasmic contact sites (MERCS), which undergo physical alterations and malfunction in AD brain. Despite ongoing research, the understanding of MERCS in AD remains in its nascent stages. We postulate that these subcellular interfaces are responsible for AD progression. Neuroinflammation contributes significantly to neurodegeneration and is primarily driven by microglia, the innate immune cells in the brain. In AD, activated microglia secrete pro-inflammatory cytokines that compromise neuronal vitality. The production of these cytokines is promoted by NLRP3 inflammasome. Although inflammasome activation has been observed at MERCS, the underlying MERCS-mediated mechanisms governing regulation of inflammasome activation remain to be elucidated.

**Methods:** Primary microglia were isolated from 3-4 months old wild-type (WT) and *App^NL-G-F^* mice (AD). MERCS ultrastructure was analyzed by transmission electron microscopy. Mitochondrial Ca^2+^ level and metabolic function were assessed using Rhod-2 AM fluorescence and Seahorse extracellular flux analysis respectively. Inflammasome activation was induced by lipopolysaccharide and nigericin and evaluated by IL-1β ELISA, caspase-1 activity assay, and ASC immunocytochemistry. MERCS were genetically modulated via siRNA-mediated knockdown of MERCS-associated proteins, and ER-to-mitochondria Ca²⁺ transfer was pharmacologically inhibited using Xestospongin C and MCU-i11. Microglial Aβ phagocytosis was quantified using fluorescence-conjugated Aβ_1-42_.

**Results:** AD microglia exhibited increased MERCS number and contact length, accompanied by a reduction in mitochondria-ER proximity. These structural changes were associated with elevated mitochondrial Ca^2+^ levels and enhanced respiratory activity, indicating metabolic reprogramming and functional change. Structural and functional decrease of microglial MERCS attenuated NLRP3 inflammasome activation and restored inflammasome-associated impairments in Aβ phagocytosis. Pharmacological inhibition of Ca^2+^ channels at MERCS identified ER-to-mitochondria Ca^2+^transfer as a key regulatory mechanism for inflammasome activation.

**Conclusions:** Our findings identify microglial MERCS remodeling as an early event in AD and establish ER–mitochondria coupling as an upstream regulator of energy metabolism, inflammation, and Aβ clearance. Targeting MERCS may therefore represent a promising strategy to modulate neuroinflammation while preserving essential microglial functions in AD.

## Background

Alzheimer’s disease (AD) is the leading cause of dementia and is a complex neurodegenerative disease characterized by progressive cognitive decline, neuronal dysfunction and loss, neuroinflammation, and the accumulation of amyloid β-peptide (Aβ) plaques and neurofibrillary tangles in the brain. Although disease-modifying anti-amyloid therapies, such as Lecanemab (LEQEMBI^®^) and Donanemab (Kisunla^®^), have recently been approved, there is currently no effective cure for AD, largely due to the intricate and multifactorial nature of its pathogenesis. Evidence suggests that the convergence of dysfunctional cellular processes, including dysregulated lipid metabolism, aberrant mitochondrial Ca^2+^ homeostasis, impaired autophagy, and deposition of Aβ plaques, compromises neuronal health and drives neurodegeneration [1, 2]. Intriguingly, these diverse processes are orchestrated at specialized subcellular interfaces known as mitochondria-endoplasmic reticulum (ER) contact sites (MERCS).

MERCS are dynamic interfaces where ER membranes and outer mitochondrial membranes are maintained in proximity (a distance ranging from ∼10 to ∼50 nm [3]) by tether protein complexes such as VAPB-PTPIP51 (Figure 1). These sites concentrate molecular machineries that coordinate essential cellular processes, such as ER-to-mitochondria Ca^2+^ transfer via the IP3R-GRP75-VDAC complex, as well as exchange of lipid and signalling molecules via other complexes [4]. ER-mitochondria coupling enable functions that are distinct from those of either organelle alone, including lipid biosynthesis, steroidogenesis, and the formation of localized mitochondrial Ca^2+^ hotspots [5–7]. In AD, catalytic components and activity of γ-secretase complexes are highly enriched at the ER side of MERCS, allowing Aβ generation at these interfaces [8, 9]. Recent proteomic analyses have revealed that MERCS harbour over 1,000 proteins, positioning these domains as central regulators of essential physiological processes, including mitochondrial metabolism, reactive oxygen species (ROS) production, inflammation, autophagy and amyloid precursor protein (APP) processing [10–13]. More importantly, multiple AD risk genes identified by a recent genome-wide association study (GWAS), such as APOE, ABCA7, TSPOAP1, APP and APH1B, localize to or are enriched at MERCS [14]. We and others have identified structural and functional alterations of MERCS as early hallmarks of AD [15–18]. Specifically, we have shown that Aβ increases MERCS formation that can affect mitochondrial functions and autophagosome biogenesis [19]. Modulation of MERCS by genetic manipulation of MERCS-associated protein can reduce Aβ production [20]. Despite these advances, MERCS research in the central nervous system largely focused on neurons or brain tissue. Alterations of MERCS in glial cells, which are critical mediators of AD progression [21, 22], remains largely unexplored.

**Figure 1.**
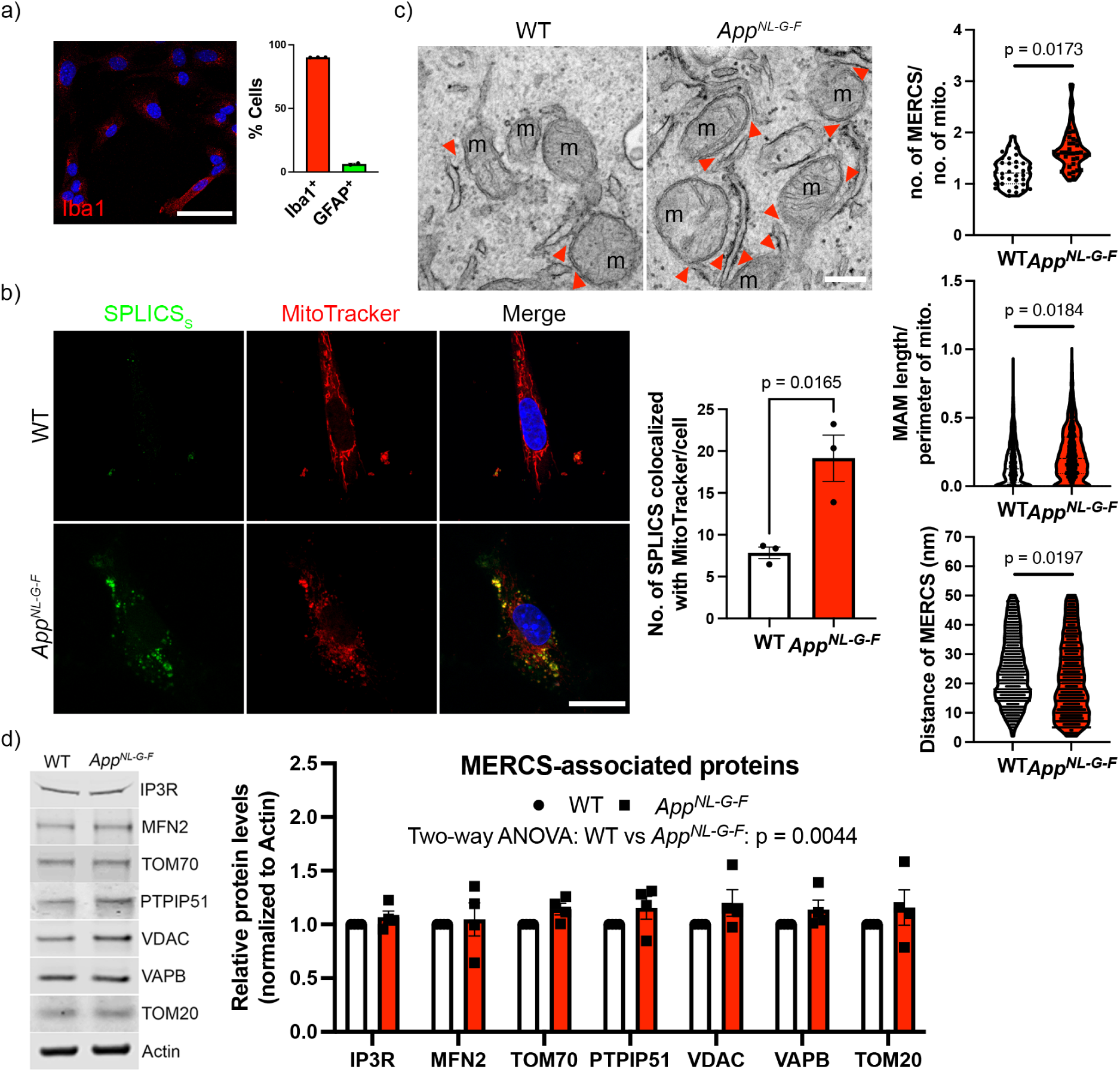
The number and length of MERCS are upregulated in primary adult *App^NL-G-F^* microglia. a) Representative image and chart showing the purity of microglial culture. Scale bar = 50 μm. n = 3. b) Representative immunocytochemistry images showing short-range mitochondria-ER contact (8-10nm; SPLICS_S_) and mitochondria (MitoTracker) and quantification of the corresponding SPLICS_S_ number (right) in primary adult microglia from wildtype (WT) and *App^NL-G-F^*(AD) mouse. Scale bar = 20 μm. Data are presented as mean ± SEM. Statistical analysis was performed using an unpaired two-tailed Student’s t-test. n = 3. c) Representative transmission electron microscopy images of mitochondria (m) and ER in WT and AD microglia illustrating MERCS. Red arrows indicate the location of MERCS. Scale bar = 200nm. Quantification of the number, the ER-mitochondria distance at MERCS, and the length of mitochondria-associated ER membrane (MAM). Data are presented as the number of MERCS per mitochondrion in individual cells, whereas the length or ER-mitochondria distance of individual MERCS are shown. Statistical analysis was performed using an unpaired two-tailed Student’s t-test on the mean from three independent experiments (n = 3). d) Representative Western blot and corresponding quantification of the levels of MERCS-associated proteins in WT and AD microglia. Data are presented as mean ± SEM. Statistical analysis was performed using two-way ANOVA followed by Šidák’s multiple-comparison test. n = 4.

Microglia are the resident immune cells of the brain and central regulators of AD progression, as evidenced by the enrichment of AD risk genes in microglia [23]. In early disease stages, they phagocytose neurotoxic Aβ aggregates providing transient neuroprotection [24]. However, defective Aβ clearance leads to sustained microglial activation and chronic neuroinflammation [25]. Activated microglia further propagate pathology by infiltrating previously unaffected brain regions and disseminating Aβ-loaded extracellular vesicles [26, 27].

A major driver of microglia-mediated neuroinflammation is the NLRP3 inflammasome [28, 29]. The canonical activation of NLRP3 inflammasome follows a two-step priming and activation process. Priming, induced by Toll-like receptor ligands like lipopolysaccharide (LPS) or Aβ, triggers NF-κB-dependent expression of NLRP3 and pro-inflammatory cytokines, such as pro-interleukin-1β. Subsequent activation is triggered by a diverse array of pathogen- or danger-associated molecular patterns (PAMPs or DAMPs) such as Aβ, mitochondrial DNA, or ROS [30]. Activation involves the assembly of NLRP3 sensor protein, ASC adaptor protein and caspase-1, leading to the proteolytic maturation of interleukin-1β (IL-1β), interleukin-18, and gasdermin D that form pores on cell membrane to release DAMPs and proinflammatory cytokines. Persistent NLRP3 inflammasome activation exacerbates neuronal damage and cognitive decline in AD [31–34], while its inhibition restores microglial Aβ phagocytosis, reduce neurotoxic cytokines, and protects against cognitive decline in AD model [28]. Emerging evidence suggests that NLRP3 inflammasome assembly occurs at MERCS in peripheral immune cells [35, 36]. This process is supported by MERCS-associated proteins such as MAVS, TXNIP and ORMDL3, which mediate NLRP3 recruitment or ER-mitochondria tethering [37–39]. However, the contribution of MERCS to inflammasome activation in microglia and the underlying mechanism remain unexplored.

Here, we provide the first evidence of significant structural remodeling of microglial MERCS in AD. We demonstrate that primary microglia derived from adult AD mouse model exhibited a significant increase in MERCS abundance, longer mitochondria-associated ER membranes and reduced ER-mitochondria distance. Mechanistically, we show that genetic modulation of MERCS via silencing the tethering protein VAPB attenuates microglial NLRP3 inflammasome activation, in part by limiting ER-to-mitochondria Ca^2+^ transfer. Importantly, MERCS modulation did not compromise microglial Aβ phagocytic capacity; instead, Aβ engulfment—normally suppressed by inflammasome activation—was partially restored. Together, our findings uncover a mechanistic link between microglial MERCS, mitochondrial Ca^2+^ signalling, and inflammasome activation. This project highlights microglial MERCS as a potential therapeutic target to mitigate neuroinflammation without impairing protective microglial functions in AD.

## Methods

### Animals

Homozygous *App^NL-G-F/NL-G-F^* (*App^NL-G-F^*) and *App^wt/wt^* (WT) mice of C57BL/6J background were housed at Karolinska Institutet animal facility with a 12-hour light/dark cycle, controlled temperature of 22-23°C, and *ad libitum* access to food and water. All animal experiments were carried out in accordance with the guidelines of the Institutional Animal Care and Use of Committee and the European Community directive (2010/63/EU) and approved by the Stockholm Animal Ethical Committee (12779-2021 & 14573-2024). Both male and female mice were used for generating primary cultures.

### Primary adult microglial culture

Primary microglial cultures were prepared from 3-4-month-old mice, according to a published protocol with minor modifications [40]. Cortices and hippocampi were dissected and preserved in ice-cold Hibernate-E (Gibco, #A1247601) for transport. Brain tissues were minced into ∼1-2 mm^3^ pieces and enzymatically dissociated in HBSS without Ca^2+^ and Mg^2+^ (5ml per brain; Gibco, #14175095) containing collagenase A (2 mg/ml; Roche, #10103586001), DNase I (14 µg/ml; Roche, #10104159001), fetal bovine serum (FBS; 250 µl per 5 ml; Gibco, #10500064), and HEPES (10 mM, Gibco). Digestion was performed at 37°C for 15 min with gentle swirling every 5 min. Following enzymatic digestion, tissues were mechanically dissociated by sequential passage through 19G, 21G and 23G needles (Terumo), triturating 8 times per needle. The resulting cell suspension was filtered through a 70 µm cell strainer into a 50 ml tube and centrifuged at 300 x g for 10 min at 4°C, max acceleration, slow deceleration. To eliminate myelin and cellular debris, the supernatant was discarded, the pellet was resuspended in 1x DPBS without Ca^2+^ and Mg^2+^ (3.1 ml per brain; Gibco) supplemented with Debris Removal Solution (0.9 ml per brain; Miltenyi Biotec, #130-109-398) and transferred to a 15ml tube. 4 ml of 1x DPBS were carefully layered on top, followed by centrifugation at 3000 x g for 10 min at 4°C, max acceleration and deceleration. After removal of the supernatant and debris layer, the remaining pellet and 2 ml leftover supernatant was brought to 14 ml with 1x DPBS and centrifuged at 1000 x g for 10 min at 4°C, max acceleration and deceleration. The pellets were resuspended in 1 ml Red Blood Cell Lysis Buffer (Roche, #11814389001) and incubated at room temperature for 5 min. Subsequently, 7 ml of 1x DPBS was added and samples were centrifuged at 300 x g for 5 min at room temperature. The final cell pellet was resuspended in DMEM/F12 supplemented with GlutaMAX (Gibco, #10565-018) and 10% FBS (medium) and plated onto non-coated 6 cm (for 1 brain) or 10 cm (for 2 brains) culture dish. Mixed glial cultures were maintained at 37°C in a humidified incubator with 5% CO₂. After 2.5-3 hours, the medium containing unattached astrocytes was removed and replaced with medium supplemented with GM-CSF and M-CSF (100 ng/ml; R&D Systems, #415-ML & 416-ML). Cultures were maintained for 7-10 days before passaging to the desired culture vessels. Cells were incubated in 0.5% Trypsin-EDTA for 10 min with dish tapping at 5 min. After detachment, cell numbers were determined using a TC20 Automated Cell Counter (Bio-Rad). Cells were used for experiments 1–2 days after reseeding. Only primary microglia at passage 1 were used for all experiments.

### Split-GFP-based contact site sensor (SPLICS) quantification

SPLICS, including the short-range (SPLICS_S_) and long-range (SPLICS_L_), kindly provided by Prof. Tito Calì (University of Padova, Italy) [41], were transfected using Lipofectamine 2000 (Invitrogen, #11668019) according to the manufacturer’s instructions. Primary microglia on 13-mm glass coverslips in 24-well plates were transfected with 0.6 µg plasmid DNA mixed with 0.6 µl Lipofectamine 2000 in 50 ul Opti-MEM (Gibco) per well and incubated for 10 min at room temperature. The transfection mixture was then added to wells containing 250 ul of culture medium. 48h after transfection, MitoTracker Deep Red (Invitrogen, #M22426) was added to the transfected cells at a final concentration of 300 nM and incubated for 30 min at 37^°^C in the presence of Hoechst 33342. Cells were then subsequently fixed with 4% paraformaldehyde (PFA) for 10 min at room temperature. After washing with PBS, coverslips were mounted using Vectashield antifade mounting medium. Fluorescence images were acquired using a 63x objective on a Zeiss LSM 980 confocal microscope and quantification of the number of SPLICS was done manually.

### Assessment of microglial bioenergetics via Oxygen Consumption Rate (OCR) and Extracellular Acidification Rate (ECAR)

Cellular respiration was assessed by measuring OCR and ECAR, which reflect mitochondrial respiration and glycolytic activity, respectively, using a Seahorse XFe96 Analyzer (Agilent Technologies). Primary microglia were seeded at a density of 1 × 10^4^ cells per well in uncoated XF96 cell culture microplates. For OCR measurements, culture medium was replaced with DMEM (pH 7.4; Sigma-Aldrich, #D5030) supplemented with 10 mM glucose, 1 mM sodium pyruvate (Sigma-Aldrich, #S8636), and 1 x GlutaMAX (Gibco). For ECAR measurements, the assay medium was prepared without glucose and pyruvate but supplemented with 1 x GlutaMAX. Prior to measurement, cells were incubated at 37°C in a non-CO₂ incubator for 15 min.

OCR was measured under basal conditions followed by sequential injections of oligomycin A (final concentration: 1 µM; Sigma-Aldrich, #75351), carbonyl cyanide-4-(trifluoromethoxy)phenylhydrazone (FCCP; final concentration: 1 µM; Sigma-Aldrich, #C2920), and a combination of rotenone (final concentration: 0.5 µM;Sigma-Aldrich, #R8875) and antimycin A (final concentration: 0.5 µM; Sigma-Aldrich, #A8674). ECAR was measured at baseline followed by sequential injections of glucose (final concentration: 15 mM), oligomycin A (final concentration: 1 µM), and 2-deoxy-D-glucose (2-DG; final concentration: 50 mM; Sigma-Aldrich; #D8375). OCR and ECAR measurements were obtained using repeated assay cycles consisting of 3 min of mixing, 0 min of waiting, and 3 min of measurement.

Data were exported using Wave software version 2.6.1 (Agilent Technologies). OCR values (pmol O_2_/min) and ECAR values (mpH/min) were normalized to total cellular protein content measured using the Pierce™ BCA Protein Assay Kits (Thermo Scientific, #23225). For OCR, basal respiration, ATP-linked respiration, maximal respiration, and spare respiratory capacity were calculated according to Seahorse XF Cell Mito Stress Test Kit User Guide. For ECAR, glycolysis, glycolytic capacity, and glycolytic reserve were calculated according to Seahorse XF Glycolysis Stress Test Kit User Guide. For each condition, OCR and ECAR values were calculated as the average of three consecutive measurements, in accordance with the manufacturer’s recommendations.

### Mitochondrial Ca^2+^ measurement using Rhod-2 AM

Mitochondrial Ca^2+^ levels were measured using mitochondria-preferring fluorescent Ca^2+^ indicator Rhod-2 AM (Invitrogen, #R1244). 0.8-1 x 10^5^ primary microglia were cultured on 35 mm glass-bottom dish (MatTek, #P35G-1.5-14-C) and washed once with pre-warmed 1 x DPBS prior to dye loading. Cells were incubated with Rhod-2 AM (1 µM) in the presence of probenecid (1 mM; MedChemExpress, #HY-B0545) and Hoechst 33342 (4 µg/ml, Invitrogen, #H3570) for 30 min at 37°C in the dark. To enable direct comparison with Seahorse OCR measurements, dye loading was performed in Seahorse OCR assay medium. Following incubation, cells were washed twice with 1 x DPBS and maintained in Seahorse OCR assay medium supplemented with 1 mM probenecid to prevent dye extrusion via organic anion transporters. Fluorescent images were acquired at 20x magnification using a Nikon CrEST X-Light V3 spinning disk confocal microscope. Rhod-2 AM fluorescence was excited at 545 nm and emission was collected between 570 and 590 nm. Mitochondrial Ca^2+^ levels were quantified as Rhod-2 fluorescence intensity per cell. Fluorescence values were normalized to WT controls and expressed as relative fluorescence intensity.

### Inflammasome induction and pharmacological inhibition

Primary microglia were seeded at a density of 0.8-1 x 10^5^ cells per well in 24-well plates containing uncoated 13-mm coverglass (Thickness no. 1.5) for subsequent IL-1β ELISA and immunocytochemistry and imaging, or at 1-2 x 10^4^ cells per well in uncoated 96-well plates. Cells were primed with 1 ng/µl lipopolysaccharide (LPS; Sigma-Aldrich, #L4391) for 3h, after which nigericin (Nig; Sigma-Aldrich, #N7143) was directly added to the LPS-containing medium to a final concentration of 3 µM and incubated for an additional 1h to induce NLRP3 inflammasome activation. Control samples received LPS treatment for 3h followed by the addition of an equivalent volume of absolute ethanol. Treatment volume was 500 µl for 24-well plates and 200 µl for 96-well plates. For pharmacological inhibition experiments, Xestospongin C (XeC; 5µM; Tocris Bioscience, #1280) or MCU-i11 (10µM; MedChemExpress, #HY-W194810) was added concomitantly with nigericin, with an incubation time of 1h.

### IL-1β ELISA

Inflammasome activation was assessed by measurement of secreted IL-1β, caspase-1 activity, and emergence of ASC speck, which was detected by immunocytochemistry (see below). Levels of secreted IL-1β in cell culture supernatants were quantified using a Mouse IL-1β ELISA Kit (Abcam, #ab197742) according to the manufacturer’s instructions. Conditioned medium was collected and centrifuged at 2000 x g for 10 min at room temperature to remove cellular debris. Clarified supernatants were diluted 1:5 prior to ELISA procedure. Absorbance was measured at 450 nm using a CLARIOstar Plus microplate reader (BMG Labtech), and original concentration of secreted IL-1β was calculated from a standard curve.

### Caspase-1 Activity Assay

Caspase-1 activity was assessed in clarified conditioned medium using the Caspase-Glo^®^ 1 Inflammasome Assay (Promega, #G9951), according to manufacturer’s instructions, as caspase-1 secretion was shown as a marker of inflammasome activation [42, 43]. 50 µl of Caspase-Glo^®^ 1 Reagent, with or without the caspase-1 inhibitor YVAD-CHO, was added to 50 µl of undiluted conditioned medium and incubated for 1h at room temperature in the dark. Luminescence was measured using a CLARIOstar Plus microplate reader. Caspase-1 activity was calculated by subtracting luminescence values obtained in the presence of YVAD-CHO from those obtained in its absence. Medium with M-CSF & GM-CSF plus Caspase-Glo^®^ 1 Reagent with or without YVAD-CHO inhibitor served as blank controls. For pharmacological inhibition experiments, to avoid interference of XeC or MCU-i11 with luminescence detection, culture medium containing these inhibitors was removed prior to the addition of 50 µl Caspase-Glo^®^ 1 Reagent, and intracellular caspase-1 activity was measured instead. Caspase-Glo^®^ 1 Reagent with or without YVAD-CHO inhibitor served as blank controls.

### Small interfering RNA (siRNA)-mediated gene knockdown

Gene knockdown was achieved by siRNA-mediated transfection. Cells were seeded in 24-well plates with 13-mm glass coverslip and allowed to adhere overnight to reach approximately 70-80% confluency at the time of transfection. Target-specific siRNAs (FlexiTube GeneSolution GS56491 for *Vapb* and GS28185 for *Tom70*; Qiagen, #1027416) and AllStars Negative Control siRNA (Qiagen, #1027280) were prepared according to the manufacturer’s instructions.

Transfection was performed using Lipofectamine 3000 (Invitrogen, #L3000001) following manufacturer’s protocol. Briefly, siRNA and Lipofectamine 3000 (0.6 µl per well) were diluted in 50 µl Opti-MEM to achieve a final siRNA concentration of 10 nM per well. The siRNA-transfection reagent complexes were incubated for 15 min at room temperature and then gently added to the centre of each well containing 250 µl culture medium. Cells were maintained under standard culture conditions and used for LPS and nigericin stimulation 24h after transfection.

### Aβ phagocytosis assay

Following LPS and nigericin stimulation, cells were washed once with 1x PBS. To assess phagocytic activity, primary microglia were incubated with Aβ_1-42_, HiLyte™ Fluor 488-labelled (5 µg/ml; Anaspec, #AS-60479-01) in 250 µl culture medium supplemented with M-CSF and GM-CSF for 2h at 37°C. Cells were then fixed for subsequent immunocytochemistry and imaging. Phagocytic uptake was quantified by measuring the total Aβ-associated fluorescence area using Fiji (ImageJ). Fluorescence values were normalized to the number of cells exhibiting positive Aβ fluorescence.

### Western blot analysis

Cells were scraped and lysed in ice-cold RIPA buffer (Thermo Scientific, #89900) supplemented with 1:100 phosphatase inhibitor cocktail (Sigma-Aldrich, #P0044) and protease inhibitor cocktail (Promega, #G6521). Lysates were transferred to 1.5 ml tubes and incubated on ice for 30 min with vortexing every 15 min. Samples were then centrifuged at 14000 x g for 5 min at 4°C, and the clarified supernatants were collected. Protein concentrations were determined using BCA assay.

Equal amounts of protein (15-20 µg) mixed with 4x Protein Sample Loading Buffer (LI-COR Biosciences, #92840004) and Amersham^TM^ ECL^TM^ Rainbow^TM^ Marker - Full range (Cytiva, #GERPN800E) were separated by SDS-PAGE using NuPAGE™ Bis-Tris Mini Protein Gels, 4–12% (Invitrogen) or NuPAGE™ Tris-Acetate Mini Protein Gels, 7% (used exclusively for IP3R detection; Invitrogen). Proteins were transferred onto Amersham™ Protran 0.45 µm nitrocellulose membrane (Cytiva). Membranes were blocked with 5% bovine serum albumin (BSA) in TBS containing 0.1% Tween 20 (TBS-T) for 1h at room temperature and incubated overnight at 4°C with primary antibodies in 5% BSA in TBS-T: IP_3_R (1:1000; Abcam, #ab5804), MFN2 (1:1000; Abcam, #ab56889), PTPIP51 (1:1000; GeneTex, #GTX54674), VDAC (1:1000; Abcam, #ab15895), TOM20 (1:1000; Santa Cruz Biotechnology, #sc-17764), TOM70 (1:1000; Santa Cruz Biotechnology, #sc-366282), and VAPB (1:1000; kindly provided by Prof. Chris Miller). Actin (1:2500; Sigma-Aldrich, #A4700) was used as a loading control. After washing with TBS-T, membranes were incubated with IRDye® 800CW Donkey Anti-Rabbit IgG (LI-COR Biosciences, #926-32213) or IRDye® 800CW Donkey anti-Mouse IgG Secondary Antibody (LI-COR Biosciences, #926-32212) for 1h at room temperature. Protein bands were visualized using an Odyssey CLx Imager (LI-COR Biosciences), and band intensities were quantified and normalized to that of actin using Fiji (ImageJ).

### Immunocytochemistry (ICC) and fluorescent microscopy

Cells cultured on glass coverslips were fixed with 4% PFA for 10 min at room temperature, permeabilized with 0.2% Triton X-100 for 5 min, and blocked with 3% BSA in PBS. Cells were incubated overnight with primary antibodies in 3% BSA at 4°C against ASC (1:200; Adipogen, #AG-25B-0006-C100), Iba1 (1:500, FUJIFILM Wako, # 019-19741), calnexin (1:100; Invitrogen, #MA3-027), TOM20 (1:100; Abcam, #ab289670). After washing, cells were incubated with appropriate fluorophore-conjugated secondary antibodies (1:500; Invitrogen).

Nuclei were counterstained with Hoechst 33342 (invitrogen). Coverslips were mounted onto glass slides using Vectashield antifade mounting medium. Fluorescence images were acquired using a Zeiss LSM 980 confocal microscope with identical acquisition settings across experimental groups. Colocalization of calnexin and TOM20 was imaged using 40x objective, while other experiments were performed using a 20x objective. Colocalization between calnexin and TOM20 was quantified using the JaCoP plugin in Fiji (ImageJ) by calculating Pearson’s correlation coefficient and Manders’ colocalization coefficient. In addition, line profile analyses were performed in Fiji (ImageJ) by drawing line regions of interest across ASC specks and extracting fluorescence intensity values to assess the spatial distribution and overlap of fluorescent signals. The area of ASC specks in images was quantified using Fiji (ImageJ).

### Transmission Electron Microscopy (TEM) and analysis

Primary microglia were detached by trypsinization and collected by centrifugation at 300 x g for 5 min. Cells were resuspended and fixed in 2.5% glutaraldehyde buffered with 0.1 M Sorensen phosphate buffer (pH 7.4) for 1h at room temperature, followed by storage at 4°C. Fixed cells were rinsed twice 0.1 M phosphate buffer and post-fixed in 2% osmium tetroxide in 0.1 M phosphate buffer (pH 7.4) 2h at 4°C. Samples underwent stepwise dehydration in a graded ethanol series and *en bloc* staining with 0.5% uranyl acetate (UAc). Following dehydrated in acetone, samples were infiltrated and embedded in LX-112 epoxy resin (Ladd Research). Ultrathin sections (∼80-100 nm) were prepared using an EM UC7 ultramicrotome (Leica Microsystems) and mounted on Formvar-coated slot grids. Sections were contrasted with UAc followed by lead citrate before examination. Imaging was performed using a HT7700 transmission electron microscope (Hitachi High-Technologies) operated at 80 kV. Digital images were acquired using a 4MPx Veleta CCD camera (EMSIS GmbH) at 49,000 x.

All mitochondria from at least 10 different cells were imaged per genotype. Number of MERCS, distance between mitochondria and ER at MERCS, length of mitochondria-associated ER membrane, mitochondrial area, perimeter, and aspect ratio, were quantified using the freehand line tool in Fiji (ImageJ). The number of MERCS per mitochondria was obtained by dividing the number of MERCS per number of mitochondria in each cell. Inter-organelle distance ≤ 50 nm between ER and mitochondria were considered MERCS. The aspect ratio was calculated by dividing the longer axis of the mitochondria profile by the shorter axis of the mitochondria profile.

### Statistical Analysis

All data are presented as mean ± standard error of the mean (SEM). Data were obtained from at least three independent experiments. *n* denotes the number of independent biological replicates. Normality was assessed using the Shapiro–Wilk test. Statistical analyses were performed using GraphPad Prism 10. Comparisons between two groups were performed using an unpaired two-tailed Student’s t-test. For TEM analysis, statistical testing was conducted using three mean values derived from the corresponding three independent biological replicates. Although individual data points representing single cells or MERCS are shown in the graphs, statistical analyses were based on replicate means rather than individual measurement. Comparisons among multiple groups were performed using one-way or two-way analysis of variance (ANOVA) followed by Tukey’s post hoc or Šidák’s multiple-comparison test respectively. The specific statistical tests used for each experiment are indicated in the figure legends. A p-value of < 0.05 was considered statistically significant.

## Results

### Microglial proximal MERCS are selectively increased in early-stage Alzheimer’s disease

To determine whether MERCS are altered in microglia during early-stage AD, we first established primary microglial cultures from the cortex and hippocampus of 3-4-month-old *App^NL-G-F^* (AD) and wild type (WT) mice. The *App^NL-G-F^*knock-in model expresses humanized APP and displays progressive Aβ plaque deposition, glial reactivity and neuroinflammation without the confounding effects of APP overexpression [44]. Cultures were highly enriched for microglia, with over 90% of cells expressing the microglial marker Iba1 (Figure 1a).

MERCS abundance was first assessed using SPLICS, which emits fluorescence when the ER and mitochondria are in close apposition [41]. Primary microglia expressing SPLICS_S_, which selectively labels proximal ER–mitochondria contacts separated by 8-10 nm, displayed a marked increase in SPLICS_S_ GFP puncta in AD compared to WT controls (Figure 1b), indicating enrichment of proximal MERCS. In contrast, labelling of distal contacts (40-50 nm) using SPLICS_L_ revealed no significant difference between AD and WT microglia (Supplementary Figure 1). These findings suggest that early-stage AD is associated with a selective increase in proximal, but not distal, MERCS in microglia.

We next validated these observations at ultrastructural resolution using TEM. ER-mitochondria (ER-MT) contacts were defined as regions with an inter-organelle distance of ≤50 nm. To account for potential alterations in mitochondrial content or morphology, the number of MERCS was normalized to mitochondrial number, and contact length was normalized to mitochondrial perimeter. AD microglia exhibited a significant increase in both the number of MERCS per mitochondrion and the length of mitochondria-associated ER membranes compared with WT controls (Figure 1c), despite no detectable difference in total mitochondrial number and area (Supplementary Figure 1). Distance-based analysis further revealed a redistribution of ER-MT distance of MERCS in AD microglia. Consistent with the SPLICS results, contacts with an ER–mitochondria distance of ≤10 nm were substantially enriched in AD microglia, whereas contacts within the 15-25 nm range were reduced, and the proportion of distal contacts (40-50 nm) remained unchanged (Figure 1c, Supplementary Figure 2). These results suggested enhanced physical coupling and possibly exchange of molecules between the two organelles.

To explore whether these structural alterations were accompanied by molecular changes, we examined the expression of key MERCS-associated tethering and functional proteins. While individual protein levels were not significantly altered, two-way ANOVA revealed a significant genotype effect when MERCS-associated proteins were analyzed collectively, indicating an overall increase in the MERCS protein in AD microglia (Figure 1d). Together, these data demonstrate that early-stage AD is characterized by a selective expansion of proximal MERCS and increased ER-mitochondria interface length in microglia, partly supported by coordinated changes in MERCS-associated proteins.

### Enhanced mitochondrial Ca^2+^ accumulation and metabolic activity in early-stage AD microglia

The physical coupling between the ER and mitochondria facilitates the bidirectional exchange of signaling molecules, including Ca^2+^ ions and lipids, and metabolic coordination between organelles. Given the increased MERCS abundance in AD microglia, we first examined mitochondrial Ca²⁺ levels using Ca²⁺ indicator Rhod-2 AM accumulating in the mitochondrial matrix. Quantitative fluorescence analysis revealed a significant increase in Rhod-2 fluorescence in AD microglia compared with WT controls (Figure 2a), suggesting elevated ER-to-mitochondria Ca^2+^ transfer.

**Figure 2.**
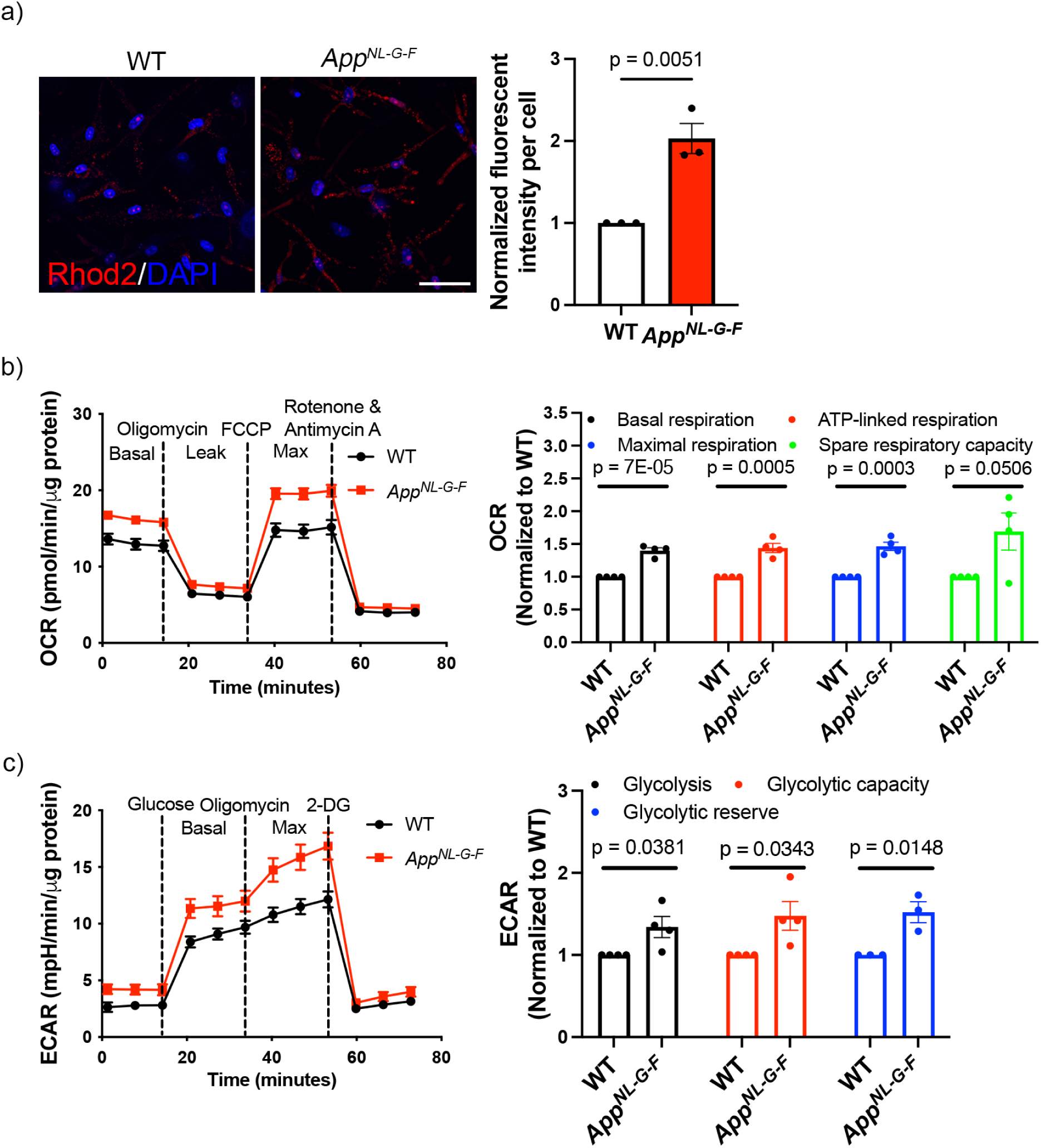
Mitochondrial Ca^2+^ levels and metabolic activity are enhanced in primary adult *App^NL-G-F^* microglia. a) Representative fluorescence images showing Rhod-2 fluorescence intensity in primary wildtype (WT) and *App^NL-G-F^* (AD) microglia loaded with the mitochondrial Ca^2+^ indicator Rhod-2 AM. Scale bar = 50 μm. Quantification showing relative Rhod-2 fluorescence intensity per cell normalized to WT levels. Data are presented as mean ± SEM. Statistical analysis was performed using an unpaired two-tailed Student’s t-test. n = 3. b) Oxygen consumption rate (OCR) and c) extracellular acidification rate (ECAR) of primary WT and AD microglia measured by Seahorse analysis. Data were collected from at least 5 wells of WT and AD microglia. OCR and ECAR values are normalized to total protein content per well. Data are presented as mean ± SEM. Statistical analysis was performed using an unpaired two-tailed Student’s t-test. n = 4.

We next assessed whether elevated mitochondrial Ca^2+^ levels were associated with altered mitochondrial function. Mitochondrial respiration was evaluated using Seahorse extracellular flux analysis under basal conditions followed by sequential perturbation of the mitochondrial electron transport chain. AD microglia exhibited higher basal oxygen consumption relative to WT controls (Figure 2b). Upon uncoupling of the proton gradient with FCCP, maximal respiration and spare respiratory capacity were also increased in AD microglia. Inhibition of ATP synthase with oligomycin further revealed enhanced ATP-linked respiration in AD microglia. These data indicate that mitochondrial respiration is augmented in microglia from early-stage AD mice.

Because late-stage AD exhibits heightened glycolysis and impaired mitochondrial respiration, we asked whether the increased mitochondrial respiration in early-stage AD microglia is also associated with reduced glycolytic metabolism. On the contrary, extracellular acidification rate (ECAR) measurements instead revealed that AD microglia displayed elevated glycolytic activity following glucose addition (Figure 2c). Inhibition of mitochondrial ATP production with oligomycin further uncovered increased glycolytic capacity and glycolytic reserve in AD microglia. Thus, early-stage AD microglia exhibit a coordinated enhancement of both mitochondrial and glycolytic metabolism, suggesting metabolic reprogramming toward increased cellular activity.

Additionally, to determine whether these functional changes were accompanied by altered mitochondrial dynamics, which can be regulated by MERCS [45–48], we quantified mitochondrial size and aspect ratio from TEM images, parameters indicative of mitochondrial fission-fusion balance. No differences were detected between AD and WT microglia (Supplementary Figure 2), suggesting that mitochondrial morphology and dynamics are preserved at this disease stage. Collectively, these findings link increased ER-mitochondria coupling to elevated mitochondrial Ca²⁺ accumulation and enhanced metabolic activity in AD microglia. Notably, these changes occur under basal conditions and in the absence of detectable alterations in mitochondrial morphology, suggesting that altered ER–mitochondria coupling, rather than changes in mitochondrial mass or dynamics, is sufficient to drive functional remodeling of microglia in early-stage AD.

### MERCS serve as platforms for NLRP3 inflammasome assembly in microglia

Microglia are central mediators of innate immune responses in the brain and contribute substantially to neuroinflammation in AD. Activation of NLRP3 inflammasome has been documented in microglia in AD patient brains and experimental models, where it exacerbates neurological decline [35]. Although MERCS have been implicated as sites of NLRP3 inflammasome assembly in other immune cell types, their role in microglia has not been clearly defined.

To address this, we first validated that adult primary microglial cultures are capable of robust NLRP3 inflammasome activation. WT microglia were primed with lipopolysaccharide (LPS) and subsequently stimulated with nigericin (LPS + Nig), a well-established inducer of NLRP3 inflammasome activation [49, 50]. Inflammasome activation was confirmed by increased IL-1β secretion (Figure 3a), elevated extracellular caspase-1 activity (Figure 3b), and the formation of ASC specks in a subset of microglia (Figure 3c). The percentage of ASC speck-positive cells in our cultures was consistent with previous reports in other immune cell types [51–53].

**Figure 3.**
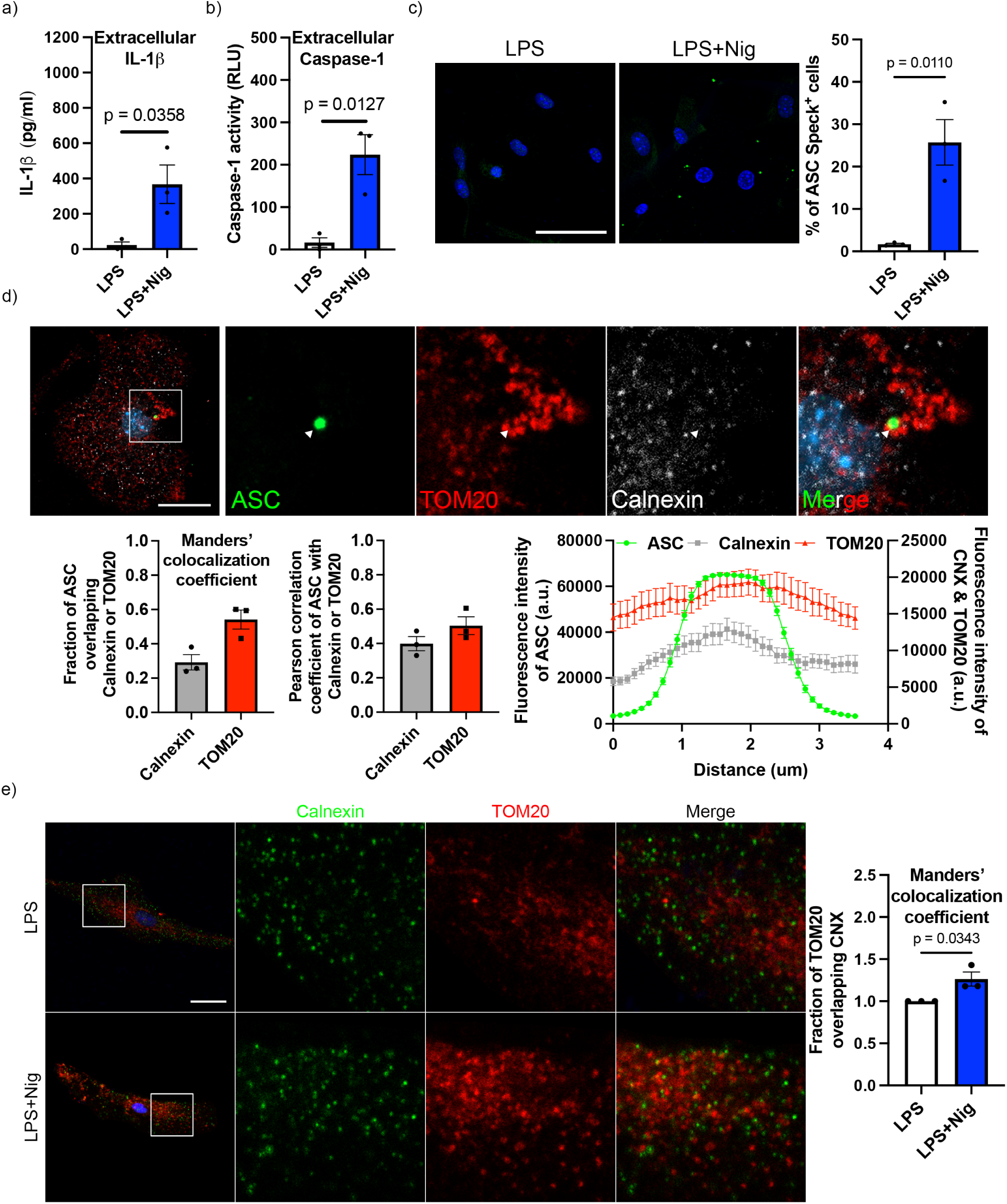
Inflammasome activation is associated with ER-mitochondria coupling in primary adult microglia. a) IL-1β levels measured by ELISA and b) extracellular caspase-1 activity of primary microglia treated with LPS for 4h or LPS for 4h with nigericin added during the final 1h (LPS + Nig). Data are presented as mean ± SEM. Statistical analysis was performed using an unpaired two-tailed Student’s t-test. n = 3. c) Representative fluorescence images of ASC immunostaining (green) in microglia treated with LPS + Nig or LPS alone. Scale bar = 50 μm. Quantification shows the percentage of ASC speck-positive cells. Data are presented as mean ± SEM. Statistical analysis was performed using an unpaired two-tailed Student’s t-test. n = 3. d) Representative fluorescence images and colocalization analysis of ASC with the ER membrane marker calnexin and the mitochondrial outer membrane marker TOM20 following LPS + Nig treatment. Scale bar = 20 μm. Manders’ colocalization coefficients indicate the fraction of ASC overlapping with calnexin or TOM20. Pearson correlation coefficients demonstrate correlation between ASC fluorescence and calnexin or TOM20. Intensity profiles across diameters of ASC specks illustrate spatial overlap. White arrows indicate the location of an ASC speck. At least 15 cells were analyzed per biological replicate. Data are presented as mean of all individual cells from independent biological triplicate ± SEM. n = 3. e) Representative images of calnexin and TOM20 in microglia treated with LPS + Nig or LPS alone. Manders’ colocalization coefficient shows the fraction of TOM20 overlapping with calnexin. Scale bar = 20 μm. Data are presented as mean ± SEM. Statistical analysis was performed using an unpaired two-tailed Student’s t-test. n = 3.

We next examined the subcellular localization of inflammasome assembly relative to MERCS using triple-label immunofluorescence for the inflammasome adaptor protein ASC, the ER membrane marker calnexin, and the outer mitochondrial membrane marker TOM20. Confocal imaging revealed that ASC specks formed upon LPS + Nig stimulation localized in close proximity to ER-mitochondria interfaces (Figure 3d). Manders’ colocalization analysis showed overlap of ASC specks with both ER membrane and outer mitochondrial membrane. Consistently, Pearson correlation analysis demonstrated a positive correlation between ASC fluorescent intensity and both calnexin and TOM20 signals. Fluorescence intensity profiling across the diameter of ASC specks further revealed enrichment of ER and mitochondrial signals at the centre of the specks, indicative of a condensed ER-mitochondria interface at sites of inflammasome assembly. These observations support the notion that NLRP3 inflammasome assembly occurs at MERCS in activated microglia.

Given this spatial relationship, we next examined whether inflammasome activation is accompanied by alterations in MERCS. Interestingly, we found that LPS + Nig stimulation significantly increased ER-mitochondria colocalization in microglia compared with LPS priming alone (Figure 3e), indicating an expansion of MERCS during inflammasome activation. Notably, an increase in MERCS abundance was also observed in AD microglia using TEM (Figure 1b & c). Together, these data identify MERCS as dynamic platforms closely associated with NLRP3 inflammasome assembly in microglia. This suggests that microglial MERCS accommodate the molecular machinery required for NLRP3 inflammasome assembly.

### Modulation of microglial MERCS attenuates NLRP3 inflammasome activation

Given that microglia in AD brains exhibit inflammasome activity, that AD microglia display increased MERCS abundance, and that MERCS are closely associated with inflammasome assembly, we next asked whether reducing MERCS could attenuate inflammasome activation in microglia. To directly test this, MERCS were genetically modulated by targeting two MERCS-associated proteins with distinct roles in ER-mitochondria coupling and signalling. Knockdown of the ER-resident tethering protein VAPB was previously shown to significantly reduce ER–mitochondria association and ER-to-mitochondria Ca^2+^ transfer [54]. In contrast, we have reported that the mitochondrial outer membrane protein TOM70 is associated with the IP3R-GRP75-VDAC complex and supports ER-to-mitochondria Ca²⁺ transfer without directly contribute to contact site abundance [55]. We first assessed the efficiency of knockdown (Supplementary Figure 3a) and confirmed that knockdown of MERCS-associated proteins effectively modulated ER–mitochondria contacts in microglia. Indeed, *Vapb* knockdown (*Vapb* KD) significantly reduced ER-mitochondria association by about two-third (Figure 4a). TOM70 depletion (*Tom70* KD) showed a non-significant trend towards reduced ER-mitochondria association, consistent with our previous report [55]. Importantly, modulation of MERCS did not affect microglia survival over the experimental time frame, indicating that subsequent experimental measurements were likely not due to the decrease of cell number (Supplementary Figure 3b).

**Figure 4.**
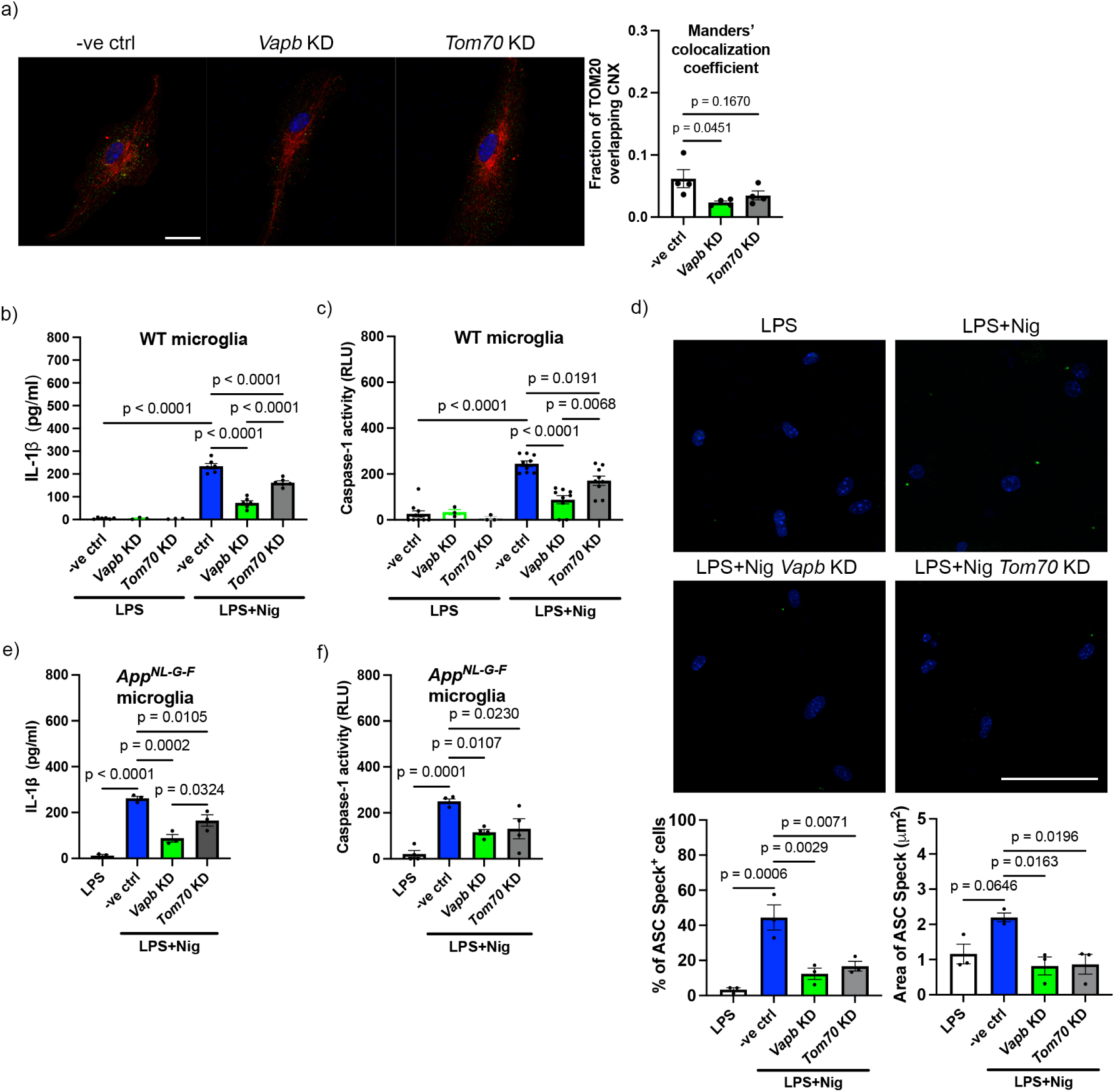
Microglial MERCS govern inflammasome assembly. a) Representative immunocytochemistry images and the corresponding analysis of Manders’ colocalization coefficient showing colocalization of calnexin and TOM20 in microglia following *Vapb* or *Tom70* knockdown. Scale bar = 20 μm. Data are presented as mean ± SEM. Statistical analysis was performed using one-way ANOVA followed by Turkey’s multiple comparisons test. n = 3. b) IL-1β levels measured by ELISA (n = 6) and c) extracellular caspase-1 activity (n = 9) of MERCS-modulated microglia with or without inflammasome induction. Data are presented as mean ± SEM. Statistical analysis was performed using one-way ANOVA followed by Turkey’s multiple comparisons test. d) Representative fluorescence images of ASC immunostaining (green) in MERCS-modulated microglia with or without inflammasome induction. Scale bar = 50 μm. Quantification shows the percentage of ASC speck-positive cells and the area of ASC specks. Data are presented as mean ± SEM. Statistical analysis was performed using one-way ANOVA followed by Turkey’s multiple comparisons test. n = 3. e) IL-1β levels measured by ELISA (n = 3) and f) extracellular caspase-1 activity (n = 4) in MERCS-modulated AD microglia with or without inflammasome induction. Data are presented as mean ± SEM. Statistical analysis was performed using one-way ANOVA followed by Turkey’s multiple comparisons test.

We then assessed the impact of MERCS modulation on NLRP3 inflammasome activation. Inflammasome activation was induced in MERCS-modulated microglia using LPS + Nig. As expected, *Vapb* KD resulted in a pronounced reduction in IL-1β secretion following LPS + Nig stimulation compared with LPS-primed cells without MERCS modulation (Figure 4b). Similarly, *Tom70* KD also decreased IL-1β release upon LPS + Nig stimulation, although to a lesser extent than *Vapb* KD. A comparable pattern was observed for caspase-1 activity, with *Vapb* KD exerting a stronger inhibitory effect than *Tom70* KD (Figure 4c). Notably, MERCS modulation in LPS-primed cells (w/o Nig) elicited only minimal inflammasome activation, indicating that reduced ER-mitochondria coupling does not activate inflammasome. This observation agrees with our findings that increased MERCS abundance is associated with inflammasome activation.

To further delineate how MERCS modulation influences inflammasome assembly, we examined ASC speck formation by immunocytochemistry. Both VAPB and TOM70 knockdown substantially reduced the proportion of inflammasome-positive microglia (Figure 4d). Strikingly, in microglia that still formed ASC specks, the average speck area in MERCS-modulated cells was reduced by approximately 50%, suggesting impaired assembly of inflammasome complexes. These findings show that modulation of MERCS attenuates NLRP3 inflammasome activation in microglia, and possibly neuroinflammation in AD. To assess the relevance of these observations in an AD context, we performed parallel experiments in primary AD microglia. Consistent with the results obtained in WT microglia, MERCS modulation in AD microglia significantly reduced caspase-1 activity and IL-1β secretion following LPS + Nig stimulation (Figure 4e & f). Collectively, these data indicate that intact MERCS structure and functions are required for complete NLRP3 inflammasome activation in both WT and AD microglia.

### ER-mitochondria Ca^2+^ signalling drives NLRP3 inflammasome activation in microglia

As knockdown of either VAPB or TOM70 – two crucial regulators of ER-to-mitochondria Ca^2+^ transfer – significantly attenuated inflammasome activation, we next tested whether Ca^2+^ signalling at MERCS is necessary for NLRP3 inflammasome activation by pharmacologically inhibiting Ca^2+^ transfer at MERCS during inflammasome activation in primary WT microglia.

Inhibition of ER Ca^2+^ release using the IP_3_R antagonist Xestospongin C (XeC) significantly reduced IL-1β secretion, an effect comparable to that observed in TOM70 KD cells stimulated with LPS + Nig (Figure 5a). Consistently, caspase-1 activity was also decreased (Figure 5b). To further confirm the role of ER-to-mitochondria Ca^2+^ transfer, we inhibited mitochondrial Ca^2+^ uptake by targeting the mitochondrial calcium uniporter (MCU) with selective inhibitor MCU-i11 [56]. Blocking mitochondrial Ca^2+^ entry similarly reduced IL-1β secretion and significantly attenuate caspase-1 activity (Figure 5c & d). Together, these findings demonstrate that efficient Ca²⁺ transfer from the ER to mitochondria is required for complete NLRP3 inflammasome activation in microglia and support a model in which MERCS-mediated Ca²⁺ signaling functions as a critical upstream regulator of inflammasome assembly and activity.

**Figure 5.**
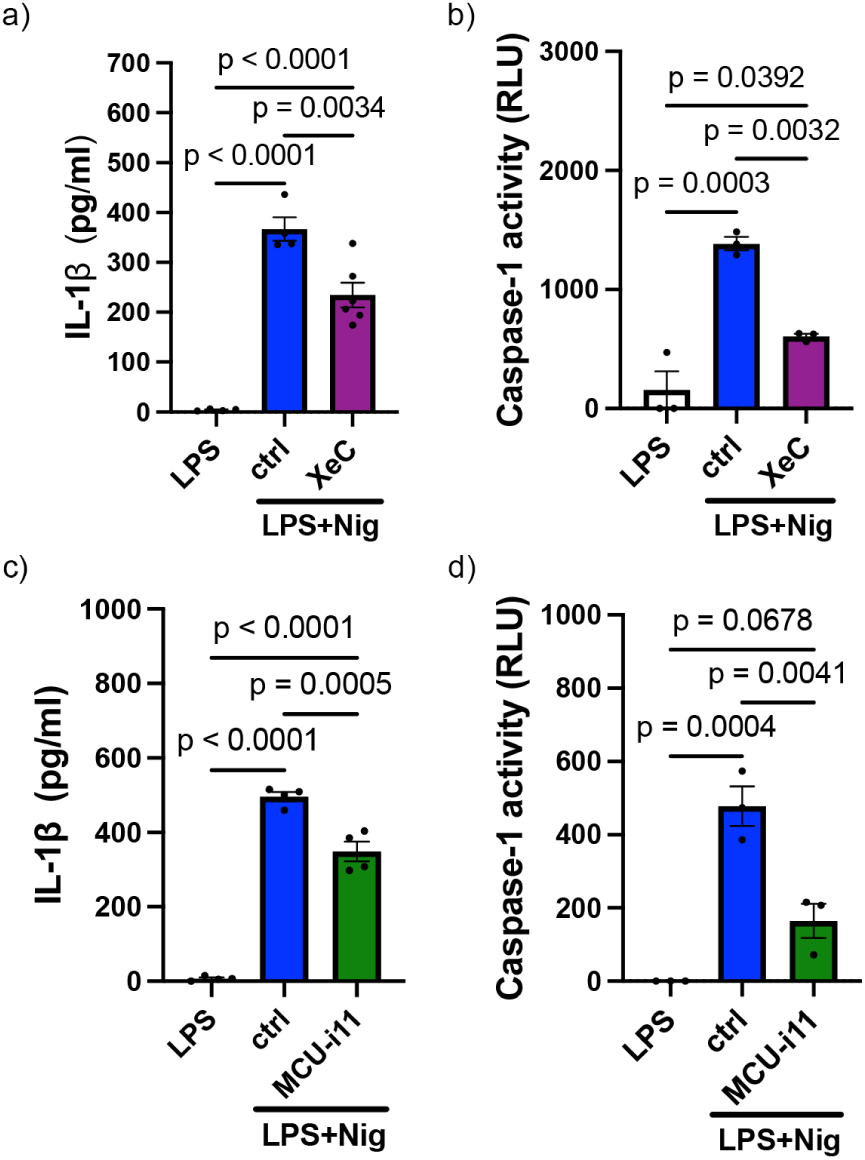
ER-to-mitochondria Ca^2+^ transfer promotes inflammasome activation in primary adult microglia. a & c) IL-1β levels and b & d) caspase-1 activity of microglia with inflammasome induction in the presence or absence of 5 μM Xestospongin C (XeC; n ≥ 3) or 10 μM MCU-i11 (n ≥ 3) at the final 1h. Data are presented as mean ± SEM. Statistical analysis was performed using one-way ANOVA followed by Turkey’s multiple comparisons test. n = 3.

### Modulation of MERCS alleviates inflammasome-associated impairment of microglial Aβ phagocytosis

In AD, activated microglia respond to Aβ by activating NLRP3 inflammasome and releasing ASC specks and pro-inflammatory mediators, including cytokines and reactive oxygen species, that can exacerbate neurotoxicity during chronic neuroinflammation [57]. In contrast, at early disease stages, microglia exert neuroprotective functions by phagocytosing and clearing Aβ. Previous studies have shown that sustained inflammasome activation and excessive inflammatory signaling impair microglial phagocytic capacity [58], representing a functional consequence of inflammasome activation.

Given our findings that MERCS modulation suppresses inflammasome activation, we wondered if reducing ER–mitochondria coupling could promote microglial Aβ phagocytosis under inflammasome-activating conditions, or perturbing MERCS would further compromise phagocytic function due to the central role of MERCS in cellular signaling. To address this, microglia with modulated MERCS were stimulated with LPS + Nig and subsequently incubated with fluorophore-conjugated Aβ to assess phagocytic activity.

Consistent with previous reports, inflammasome activation markedly reduced microglial phagocytosis, as evidenced by a significant decrease in the Aβ-engulfing microglia (phagocytic microglia) compared with LPS-primed controls (Figure 6). In parallel, the intracellular Aβ burden per phagocytic cell, quantified as the cytoplasmic Aβ-positive area, was also significantly diminished. Strikingly, modulation of MERCS through either *Vapb* or *Tom70* knockdown rescued this impairment (Figure 6). MERCS-modulated microglia subjected to inflammasome activation exhibited a restored proportion of phagocytic cells, as well as a normalized amount of internalized Aβ relative to LPS-primed controls.

**Figure 6.**
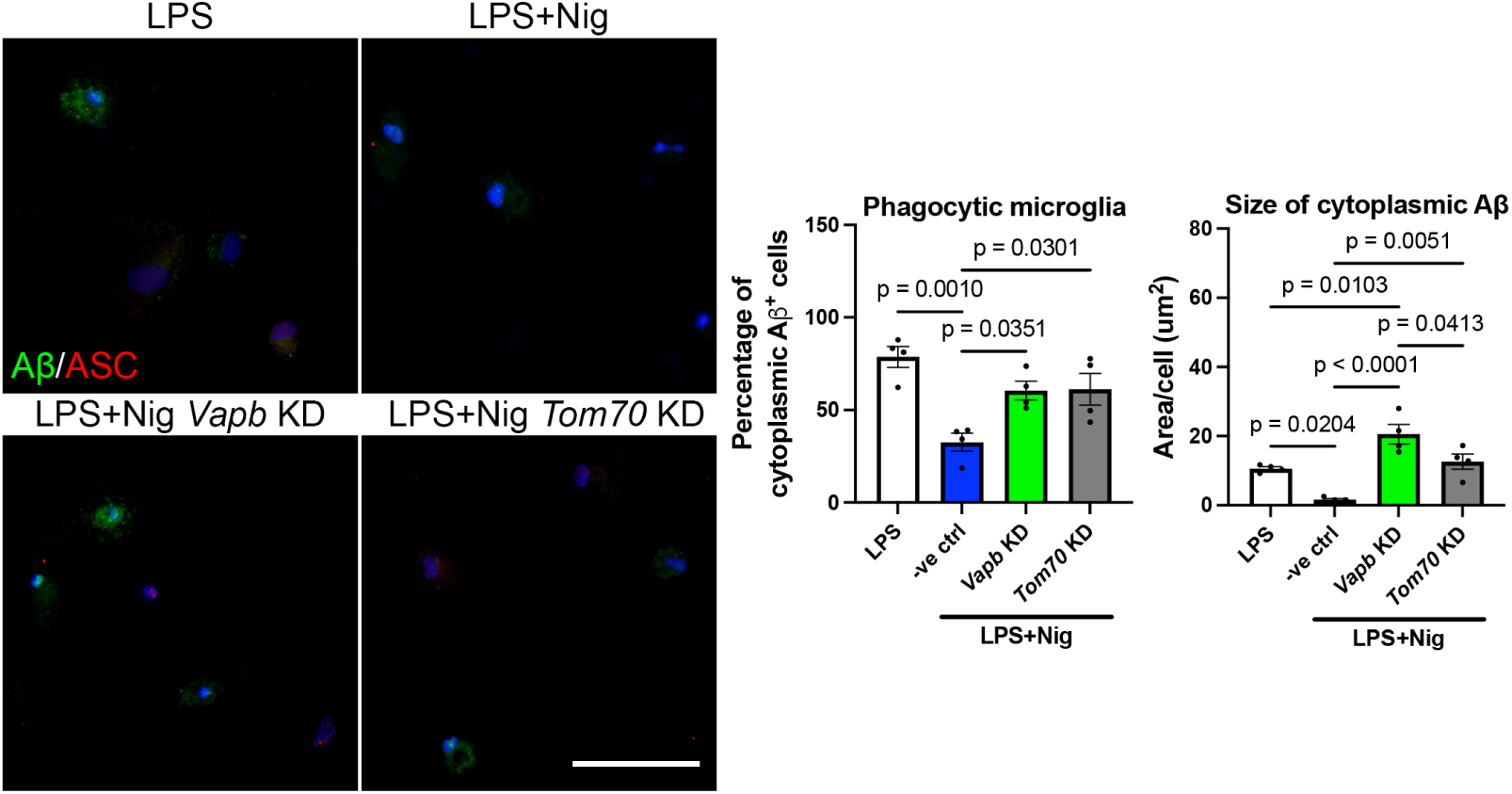
Modulation of microglial MERCS restores inflammasome-associated Aβ phagocytosis impairment. Primary adult microglia were incubated with 5 µg/ml HiLyte™ Fluor 488–conjugated Aβ₁₋₄₂ for 2h at 37°C, followed by fixation and fluorescence imaging. Representative images show intracellular Aβ uptake (green) and ASC speck. Scale bar = 50 μm. Quantification shows the percentage of phagocytic microglia, defined as cells exhibiting detectable intracellular Aβ fluorescence, and the cytoplasmic Aβ-positive area per cell. Data are presented as mean ± SEM. Statistical analysis was performed using one-way ANOVA followed by Turkey’s multiple comparisons test. n = 4.

Together, these findings indicate that excessive ER–mitochondria coupling and inflammasome activation are associated with impaired microglial phagocytosis, and that attenuation of MERCS alterations restores this essential homeostatic function. These data suggest that MERCS modulation not only suppresses excessive inflammatory responses but also helps preserve microglial clearance capacity during inflammation, highlighting MERCS as a key regulator of neuroinflammation and homeostatic microglial responses in AD.

## Discussion

The present study identifies MERCS as central regulators of microglial metabolism, inflammasome activation, and phagocytosis. Using adult primary microglia derived from the *App^NL-G-F^* knock-in mice, we demonstrate that early pathological changes are characterized by a selective expansion of proximal MERCS, accompanied by enhanced ER-to-mitochondria Ca²⁺ transfer, and upregulation of glucose metabolism. We further establish MERCS as dynamic platforms for NLRP3 inflammasome assembly and show that genetic or pharmacological modulation of MERCS-associated Ca²⁺ signaling attenuates inflammasome activation while partially restoring inflammasome-associated impairment of Aβ phagocytosis. Together, these findings provide new insight into how MERCS may govern neuroprotective and neurotoxic states of microglia during AD progression.

A major novelty of this study is the use of adult primary microglia enabling direct visualization and specific assessment of MERCS architecture independent of other brain cell types. Increased MERCS abundance in AD microglia likely reflects cell-intrinsic alterations or the lasting effect of prior *in vivo* exposure to Aβ rather than *de novo* Aβ production in culture, as AD microglia themselves generate negligible amounts of Aβ. To date, the structural and functional characterization of MERCS in microglia remains largely unexplored, particularly in AD. Notably, prior work in a chronic stress-induced depression mouse model demonstrated that reducing ER-to-mitochondria Ca^2+^ transfer via depletion of GRP75, a component of the MERCS Ca^2+^ shuttling complex, suppressed inflammasome activation and improved depressive-like behavior [59]. These findings support the role for MERCS-mediated Ca^2+^ flux in microglial inflammatory responses. Our study extends this concept to neurodegeneration.

Interestingly, the expansion of MERCS in AD microglia mirrors our previous observations in AD neurons and astrocytes [15, 16, 19, 60], suggesting that MERCS remodeling may represent a conserved cellular stress response across brain cell types. However, the functional consequences appear to be cell-type specific. In neurons, MERCS regulate APP processing and Aβ generation. In contrast, neurons lack robust inflammasome activity, whereas microglial MERCS govern inflammasome-driven cytokine release, underscoring the importance of cell-type specific investigation.

### Functional implications of proximal MERCS expansion

Our data reveal selective enrichment of short-range (≤10 nm) MERCS in early-stage AD microglia without changes in distal contacts or detectable alterations in mitochondrial morphology. This suggests that MERCS remodeling precedes overt mitochondrial fission reported in microglia in later stages of AD [61, 62].

ER-MT distance determines the function of MERCS, as distinct protein complexes for signalling events are recruited to different inter-organelle gaps [3]. Proximal contacts (∼10 nm) are thought to favour lipid exchange, whereas intermediate distances (∼20 nm) are considered optimal for Ca²⁺ transfer between ER and mitochondria [63, 64]. Interestingly, although we observed a reduction in MERCS abundance within the 15–25 nm range, mitochondrial Ca²⁺ levels were elevated in AD microglia. This apparent paradox may be explained by several non-mutually exclusive mechanisms. First, AD microglia exhibited an increased length of ER–mitochondria interface per mitochondrion, which could accommodate a greater number of Ca²⁺ transfer complexes despite altered distance distribution. Second, MERCS function is determined not only by distance but also by dynamic remodelling—features that remain underexplored [65]. Third, we observed a trend toward upregulation of Ca²⁺ transport components such as IP3R and VDAC, which may enhance ER-to-mitochondria Ca²⁺ flux. Together, these factors likely converge to elevate mitochondrial Ca²⁺ levels despite shifts in contact distance.

Enhanced mitochondrial Ca²⁺ accumulation provides a mechanistic link between MERCS expansion and the hypermetabolic phenotype of early AD microglia. Several Ca²⁺-sensitive TCA cycle enzymes, such as isocitrate dehydrogenase and ɑ-ketoglutarate dehydrogenase, could have higher activity to boost respiratory efficiency. Consistent with this, our Seahorse analyses revealed elevated basal respiration in AD microglia. These findings align with our prior reports of mitochondrial hypermetabolism in early AD brain tissue [66], which should be largely attributed to neurons and astrocytes due to their abundance in the brain. Our results suggest that this hypermetabolic state is likely not cell-type specific but instead reflects a broader mitochondrial adaptation across brain cell populations.

Concurrently, AD microglia displayed enhanced glycolytic activity, a feature observed at later disease stages, indicating early and persistent metabolic reprogramming. Mitochondrial metabolism undergoes a stage-dependent shift from hyperfunction to failure. Such metabolic plasticity may initially support protective microglial functions but later contribute to dysfunction, potentially in association with microglial senescence and chronic cytokine secretion [67–69].

### MERCS as platforms for inflammasome assembly

Our work demonstrates that MERCS serve as platforms for NLRP3 inflammasome assembly in microglia. ASC specks localized preferentially to mitochondria while maintaining close association with ER membranes, suggesting the involvement of both organelles in inflammasome activation. Genetic and pharmacological modulation of MERCS further showed that intact ER–mitochondria coupling and Ca²⁺ transfer is required for complete inflammasome activation. Knockdown of VAPB, an ER-resident tether that contributes to ER-mitochondria apposition and Ca^2+^ handling, produced stronger suppression of inflammasome activity than knockdown of TOM70, a mitochondrial protein involved in Ca²⁺ transfer without markedly altering contact abundance. Consistently, inhibition of IP3R or MCU attenuated inflammasome activation to a similar extent as TOM70 knockdown, establishing Ca²⁺ flux across MERCS as a key upstream regulator. The differential effects of VAPB and TOM70 knockdown suggest that MERCS functions beyond Ca²⁺ transfer contribute to inflammasome activation.

Mitochondria contribute to inflammasome activation through multiple mechanisms, including ROS production, Ca²⁺ influx, mitochondrial DNA release and K^+^ efflux [70]. In addition to these danger signals, specific mitochondrial proteins and lipids, such as cardiolipin, have been shown to directly facilitate inflammasome activation [37–39, 71, 72]. In parallel, ER stress can trigger inflammasome signalling through several pathways, including IRE-1ɑ-initiated mitochondrial damage [73–75]. Our findings extend the knowledge by highlighting the role of ER membranes at contact sites, where IP3R-mediated Ca²⁺ release may potentiate mitochondrial stress or activate signaling pathways promoting inflammasome activation at MERCS.

Importantly, we further demonstrate that inflammasome activation is associated with enhanced ER-mitochondria coupling. Inflammatory signaling itself has been implicated in upregulating ER-mitochondria coupling [30, 76–78], suggesting the existence of a feed-forward mechanism in which enhanced MERCS facilitate inflammasome assembly, while inflammasome signaling further remodels MERCS. NLRP3 inflammasome activation has been shown to exacerbate mitochondrial dysfunction, impair mitophagy, and sustain inflammatory signaling [79, 80]. Collectively, these observations position MERCS not as passive structural scaffolds but as dynamic regulators of inflammatory signaling in microglia.

### MERCS, phagocytosis, and microglial state transitions

Beyond inflammatory signaling, a novel aspect of this work is the identification of a functional link between MERCS and microglial phagocytosis. Phagocytosis is an energy-demanding process that requires coordinated cytoskeletal remodeling, calcium signaling, membrane trafficking, and lipid redistribution [81–83]. MERCS are positioned to regulate these processes by controlling mitochondrial ATP production, ER–mitochondria calcium transfer, and phospholipid exchange. Excessive MERCS coupling during inflammasome activation may therefore disrupt metabolic flexibility and calcium homeostasis, compromising the bioenergetic capacity and membrane dynamics required for efficient Aβ uptake and clearance.

Consistent with previous reports that chronic inflammation impairs Aβ clearance [28, 84, 85], we found that inflammasome activation markedly reduced microglial phagocytic capacity. Strikingly, MERCS modulation not only attenuated inflammasome activation but also restored both the proportion of phagocytic microglia and the amount of internalized Aβ under inflammatory conditions. These findings suggest that microglial MERCS function as a molecular switch governing the balance between protective Aβ clearance phenotype and maladaptive inflammatory states.

### Therapeutic implications and limitations

Reprogramming microglia to limit neurotoxicity while preserving protective responses has become a major focus in AD therapy. Anti-Aβ immunotherapies such as Lecanemab (Leqembi^©^) have demonstrated that enhancing microglial plaque clearance can slow disease progression, supporting the feasibility of microglial reprogramming as a therapeutic strategy [86, 87]. However, Aβ-targeting therapies may not directly address intracellular dysfunctions that contribute to disease progression. Given the heterogeneity and interconnected nature of AD pathology, targeting single pathological feature is unlikely to be sufficient. Accordingly, therapies targeting immune and inflammatory pathways constitute one of the largest categories in AD clinical trials [88–92], reflecting the recognition that resolving neuroinflammation is a critical complement to amyloid removal.

By identifying MERCS as upstream regulators of Ca²⁺ signaling, metabolism, inflammasome activation, and phagocytosis, our study highlights MERCS as promising therapeutic targets that could complement existing anti-amyloid strategies. This aligns with emerging multi-pathway strategies, such as the sigma-1 receptor agonist blarcamesine, which enhances autophagy, supports mitochondrial function, modulates ER stress responses, and dampens chronic neuroinflammation. In this context, our previous work showing that the phytochemical luteolin upregulates MERCS, boosts mitochondrial function, and ameliorates behavioral deficits in a Huntington’s disease model highlights the therapeutic potential of targeting ER-mitochondria communication [93]. Notably, while luteolin enhances MERCS in that context, our present findings indicate that reducing excessive MERCS coupling may be beneficial in AD, underscoring the need for disease- and stage-specific modulation of MERCS.

Several limitations should be acknowledged. Knockdown of VAPB or TOM70 may influence additional ER- or mitochondria-organelle contacts, as these proteins also participate in contacts beyond ER-mitochondria interfaces [94–99]. Thus, the observed effects cannot be attributed exclusively to MERCS modulation. Future studies targeting more MERCS components such as PTPIP51 on the mitochondrial side will be necessary to refine mechanistic specificity. Moreover, the acute nature of *in vitro* inflammasome induction and transient knockdown of MERCS-associated protein does not capture long-term consequences on cell viability or disease progression. Comprehensive *in vivo* studies across multiple stages of AD will therefore be required to fully define the temporal dynamics of MERCS remodeling and to identify optimal therapeutic windows.

## Conclusions

In summary, our study identifies remodeling microglial MERCS as an early event of AD in a mouse model. Enhanced ER–mitochondria coupling is associated with Ca²⁺-dependent metabolic reprogramming and supports NLRP3 inflammasome activation in microglia. Importantly, genetic modulation of MERCS attenuates inflammasome activation and restores the inflammasome-associated impairments in microglial Aβ phagocytosis, highlighting ER-to-mitochondria Ca^2+^ transfer as a key regulatory mechanism for inflammasome activation. These findings establish MERCS as functional regulators of microglia-driven inflammation and Aβ clearance and identify ER-mitochondria coupling as a critical upstream interface linking metabolism, inflammation, and phagocytic function in AD. Targeting MERCS may therefore represent a promising strategy to mitigate neuroinflammation while preserving essential microglial functions. This work supports a more comprehensive and multi-targeted approach to AD therapy.

## List of abbreviations

AD: Alzheimer’s disease
APP: Amyloid precursor protein
ASC: Apoptosis-associated speck-like protein containing a CARD
Aβ: Amyloid β-peptide
DAMPs: Damage-associated molecular patterns
ER: Endoplasmic reticulum
GRP75: 75kDa glucose-regulated protein
GWAS: Genome-wide association study
IL-1β: Interleukin-1β
IP3R: Inositol 1,4,5-trisphosphate receptor
LPS: Lipopolysaccharide
MERCS: Mitochondria-ER contact sites
Nig: Nigericin
NLRP3: NLR family, pyrin domain containing 3
PAMPs: Pathogen-associated molecular patterns
PFA: Paraformaldehyde
PTPIP51: Protein tyrosine phosphatase-interacting protein-51
ROS: Reactive oxygen species
VAPB: Vesicle-associated membrane protein-associated protein B
VDAC: Voltage-dependent anion channel
XeC: Xestospongin C

## Declarations

### Ethics approval and consent to participate Stockholm

Animal Ethical Committee *#*12779-2021 and #14573-2024.

### Consent for publication

N/A

### Availability of data and materials

The datasets used and/or analyzed during the current study are available from the corresponding author on reasonable request.

### Authors’ contributions

*MHC:* project planning, methodology, investigation, data analysis, writing—original draft preparation *MA:* conceptualization, methodology, resources, writing-review and editing, *LN:* methodology, writing-review and editing

## Acknowledgements

We are grateful to Professor Christopher Miller (Department of Basic and Clinical Neuroscience, King’s College of London, UK) for kindly providing the anti-VAPB antibodies, and Professor Tito Calì (Department of Biomedical Sciences, University of Padova, Italy) for generously sharing the SPLICS_S_ and SPLICS_L_ constructs. We would like to acknowledge the contributions from the EMil core facility at the Department of Laboratory Medicine financed by the Infrastructure Board at Karolinska Institutet for providing the electron microscopy analysis. We also thank Florian Salomons (Biomedicum Imaging Center, Karolinska Institutet) for expert microscopy assistance. We thank Dr. Noah Moruzzi for assistance with the Seahorse metabolic experiments.

