## Supplemental figures for "Modulation of mitochondria-ER contacts decrease inflammasome formation and restores amyloid β-peptide phagocytosis in adult mouse microglia"

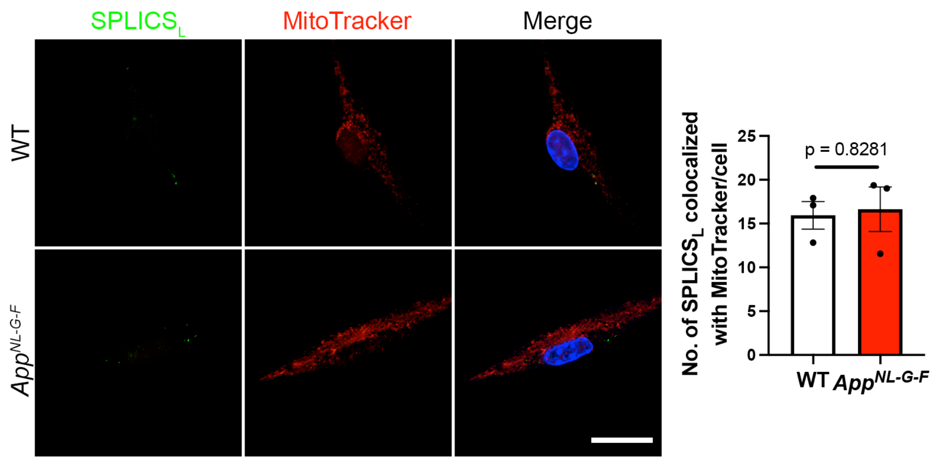


**Supplementary Figure 1. Distal MERCS are not upregulated in primary microglia from adult *App*$^{NL\text{-}G\text{-}F}$ mice.** Representative immunocytochemistry images showing long-range mitochondria-ER contact (40-50nm; SPLICS$_L$) and mitochondria (MitoTracker) and quantification of the corresponding SPLICS$_L$ number (right) in primary adult microglia from wildtype (WT) and *App*$^{NL\text{-}G\text{-}F}$(AD) mouse. Scale bar = 20 µm. Data are presented as mean ± SEM. Statistical analysis was performed using an unpaired two-tailed Student's t-test. n = 3.

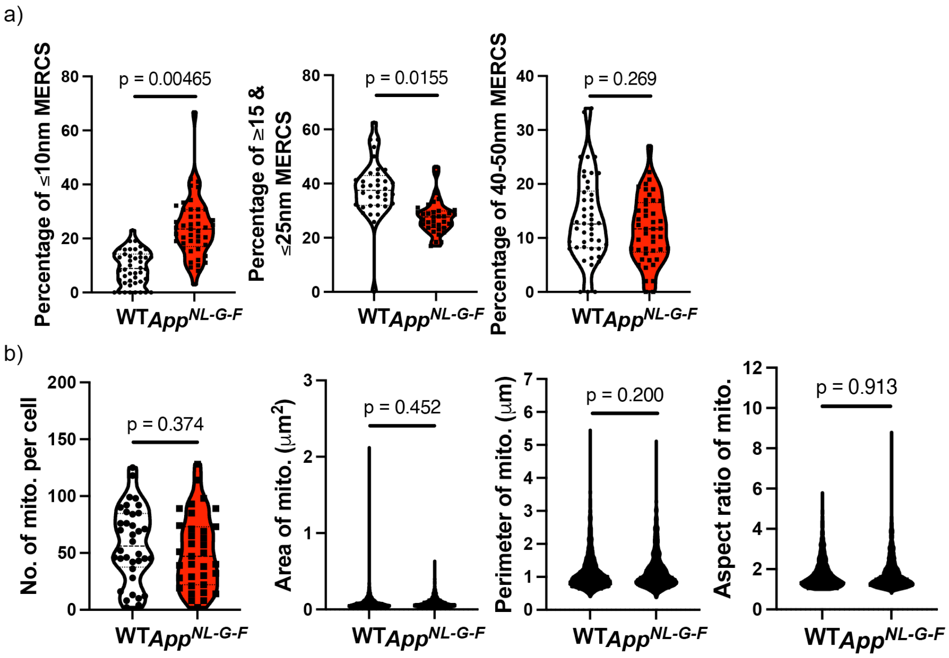


**Supplementary Figure 2. Proximal MERCS are upregulated, whereas mitochondrial morphology remains unchanged in AD microglia.** a) Quantification of the percentage of MERCS categorized by inter-organelle distance in individual cells based on transmission electron microscopy (TEM) images. Statistical analysis was performed using an unpaired two-tailed Student's t-test on the mean from three independent experiments (n = 3). b) Quantification of mitochondrial morphology parameters, including number per cell, area, perimeter, and aspect ratio, derived from TEM images. Data are presented as mitochondrial number in individual cells, whereas the area, perimeter, and aspect ratio of each mitochondrion are shown. Statistical analysis was performed using an unpaired two-tailed Student's t-test on the mean from three independent experiments (n = 3).

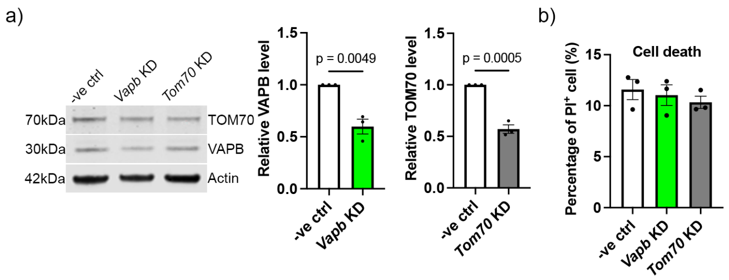


**Supplementary Figure 3. Short-term modulation of microglial MERCS does not induce cell death.** a) Representative Western blot images and quantification of VAPB and TOM70 levels in microglia following *Vapb* or *Tom70* knockdown. Data are presented as mean ± SEM. Statistical analysis was performed using unpaired two-tailed Student's t-test. n = 3. b) Quantification of propidium iodide-positive microglia following *Vapb* or *Tom70* knockdown. n = 3. Data are presented as mean ± SEM.
